# A defined 2D system for generating and expanding human basal radial glia from iPSCs

**DOI:** 10.64898/2026.04.10.714524

**Authors:** Annasara Artioli, Matteo Gasparotto, Andrea Carlo Rossetti, Yannick Hass, Anne Hoffrichter, Ryszard Wimmer, Fabio Marsoner, Raquel Perez Fernandez, Catello Guida, Philipp Koch, Alexandre D. Baffet, Ammar Jabali, Julia Ladewig

## Abstract

Basal radial glia (bRG) drive human cortical expansion but remain underrepresented *in vitro*. We established a defined and expandable 2D system for efficient generation of human bRG from iPSCs. These cells recapitulate canonical molecular signatures, hallmark somal translocation behaviours, intrinsic differentiation potential, and integration into organoid tissue. Network-based analyses identified PAK2 as a regulator of mitotic somal translocation, illustrating the system’s utility for mechanistic interrogation of bRG biology.

## Main text

Modelling human corticogenesis is constrained by limited access to fetal tissue. Although cerebral organoids have advanced developmental studies, they underrepresent basal radial glia (bRG, also known as outer RG)^1–4^. Enriched in the outer subventricular zone (oSVZ), bRG drive neocortical expansion and are defined by a characteristic molecular profile and somal translocation behaviours, including mitotic and interphase somal translocation (MST and IST)^5,6^. bRG dysregulation contributes to neurodevelopmental disorders (NDD)^7,8^, and bRG-like transcriptional states are reactivated in glioblastoma, promoting tumor heterogeneity and invasion^9^. However, current approaches allow only limited expansion of bRG from primary tissue or organoids^10^, restricting scalability of the analyses. Reliable, expandable, and experimentally accessible human bRG models are therefore essential to dissect corticogenesis and its perturbation in disease.

Here, we establish a scalable 2D-culture system enabling on-demand generation of human bRG from induced pluripotent stem cells (iPSC) (**Fig. 1A**). Building on our previously described RG differentiation paradigm, iPSCs were first patterned into neuroepithelial (NE) cells (Stage-1)^2^, followed by differentiation into RG (Stage-2)^11^. A subset of Stage-2 RG exhibited bRG-associated features (**Supp. Fig. 1A-B**). To stabilize and enrich the bRG state, we performed a targeted growth-factor screen guided by pathways implicated in oSVZ progenitor maintenance^12^ (**Supp. Fig. 2A-B**). Several conditions increased proliferation. However, YAP1 activation through TRULI^13^ and EGF exposure^12^ compromised long-term stability (**Supp. Fig. 2C**), while SHH activation and LIF signalling were excluded due to their potential to alter regional identity^14^ or lineage output^15^. Among the tested conditions, PTN and PDGF-D reproducibly increased pVIM^+^ cells and Nestin^+^ processes length. BDNF pathway activation was included as a neurotrophic factor to support culture expansion and growth (**Sup. Fig. 2C-E**). This niche-inspired combination yielded a stable Stage-3 population expandable for >15 passages.

**Figure 1.**
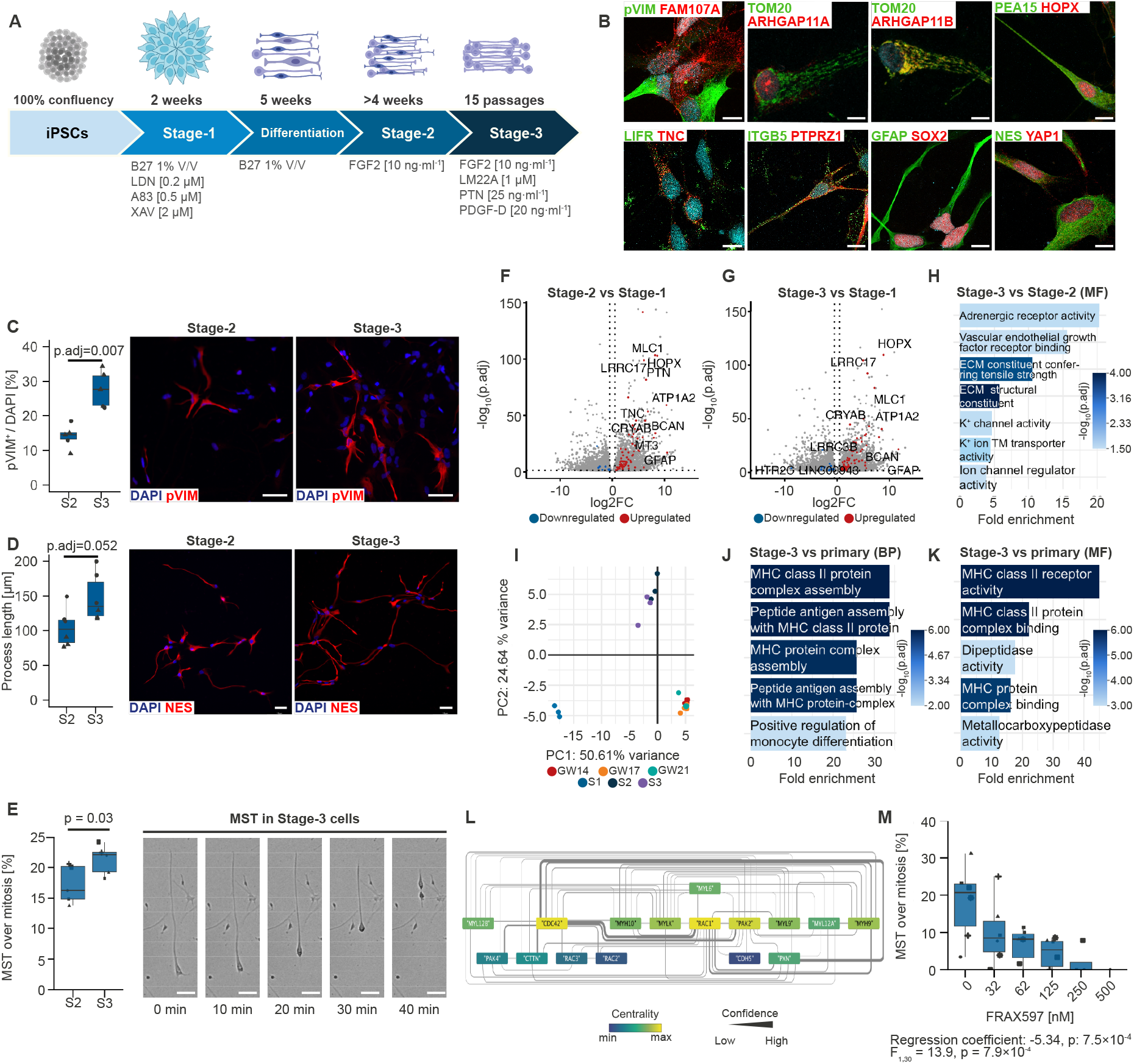
Generation and molecular-functional characterization of human iPSC-derived bRG in 2D culture. **A**. Schematic overview of the differentiation protocol. **B**. Immunostaining of Stage-3 bRG showing expression of bRG-associated marker combination. Scale bars 10 µm. **C-E**. Quantification and imaging of pVIM^+^ mitoses (C), Nestin^+^ process length (D) and MST (E) in Stage-2 and Stage-3 cultures. Scale bars 50 µm. **F-G**. Differential gene expression across differentiation stages. **H**. GO-enrichment of Stage-3 versus Stage-2. **I-K** Comparison between *in vitro* and primary bRG. **L**. Protein-protein interaction network of candidate MST regulators. **M**. Effect of PAK2 inhibition on MST frequency.

Stage-3 cultures exhibited bRG identity, expressing a characteristic marker combination including TNC, PTPRZ1, FAM107A, HOPX, LIFR and ITGB5^3,16–18^, together with nuclear YAP1 localization^19^ (**Fig. 1B, Supp. Fig. 1**). HuC/D^+^ neurons were rare and TBR2^+^ intermediate progenitors were absent (**Supp. Fig. 1**). Compared to Stage-2, Stage-3 cultures showed increased pVIM^+^ mitoses, elongated Nestin^+^ processes, and higher MST frequency (**Fig. 1C– E**). Stage-3 cultures also exhibited IST (**Supp. Fig. 2F**), supporting acquisition of hallmark bRG behaviours in vitro.

To benchmark transcriptional identity, we curated a set of 111 bRG-associated genes (**Supp. Table S1**) and performed bulk RNA-seq across Stage-1 NE, Stage-2 RG, Stage-3 and primary fetal bRG. Stage-2 and Stage-3 upregulated canonical RG markers relative to Stage-1, (**Fig. 1F-G, Supp. Tables S2-S4**). Compared to Stage-2, Stage-3 showed enrichment of ECM and membrane potential-related functions, reduced apical junction/surface programs and metabolism-related transcriptional signatures (**Fig. 1H, Supp. Figure 2G; Supp. Tables S5-S7**)^15,20^, consistent with a transition towards an oSVZ-like, dispersive bRG state. Principal component analysis placed Stage-2 and Stage-3 close to primary bRG and distinct from Stage-1 (**Fig. 1I**). Differences between Stage-3 and primary bRG are largely restricted to immune/MHC class II categories, consistent with the absence of immune signals in vitro, whereas remaining signatures were highly similar (**Fig. 1J-K, Supp. Table S8**).

To demonstrate mechanistic utility, we focused on MST, a defining yet incompletely characterized feature of bRG. Literature-guided interaction network analysis identified PAK2, a CDC42-regulated kinase involved in actin and microtubule dynamics^21^, as a top candidate to modulate MST (**Fig. 1L**). Pharmacological inhibition of PAK2^22^ reduced MST frequency in a dose-dependent manner without affecting overall cell motility (**Fig. 1M, Supp. Fig 2H**) establishing PAK2 as a regulator of MST in bRG *in vitro*.

To resolve cellular heterogeneity and lineage dynamics, we performed single-nuclei RNA sequencing (snRNA-seq) on Stage-3 cultures maintained without passaging for extended periods, as well as after three months of growth factor withdrawal. Unsupervised clustering identified seven populations (**Fig. 2A-C**), including canonical bRG-I and bRG-II states. bRG-II was enriched for cytoskeletal and motility-associated genes (*MYL12A/B, CDC42, PAK2*), consistent with its role in dynamic bRG behaviours, and showed increased metabolic and transcriptional programs relative to bRG-I (**Fig. 2D, Supp. Table S9**). An ECM-enriched RG population (ECM-RG) was also detected, and cell-cell interaction analyses highlighted ECM-associated signalling (**Fig. 2E-F**). Reference-based mapping onto a primary human fetal cortex atlas positioned progenitor and neuronal populations predominantly within post-GW20 stages (**Supp. Fig. 3A**), a developmental window associated with increased oSVZ interneuron output^23^. Consistent with this temporal annotation, growth factor withdrawal condition revealed an interneuron progenitor cluster^15^ (INP; *GAD2, DLX5, BEST3*) and downstream cortical interneurons (cIN; *DLX5/6, GAD1/2, FOXG1, ARX*) largely lacking ganglionic eminence (GE) markers (*LHX6, NKX2-1, LMO1*)^23–25^, and a migratory interneuron population (mIN; *DCC, NRCAM, SLIT2*; **Fig. 2C, Supp. Fig 3B**). Pseudotime analysis supports progression from bRG-I through INP towards cIN stages (**Fig. 2G**). Immunostainings of directed neuronal differentiated Stage-3 cultures similarly preferentially generated DLX5^+^/GAD1^+^/GAD2^+^/GABA^+^ interneurons, whereas projection neuron markers (BRN2, *SATB2, TBR1*) were rare and *NKX2-1* remained absent (**Fig. 2H**). Directed astrocyte differentiation induced GFAP^+^ cells with astroglial morphology, indicating retained glial lineage competence (**Supp. Fig. 3C**).

**Figure 2.**
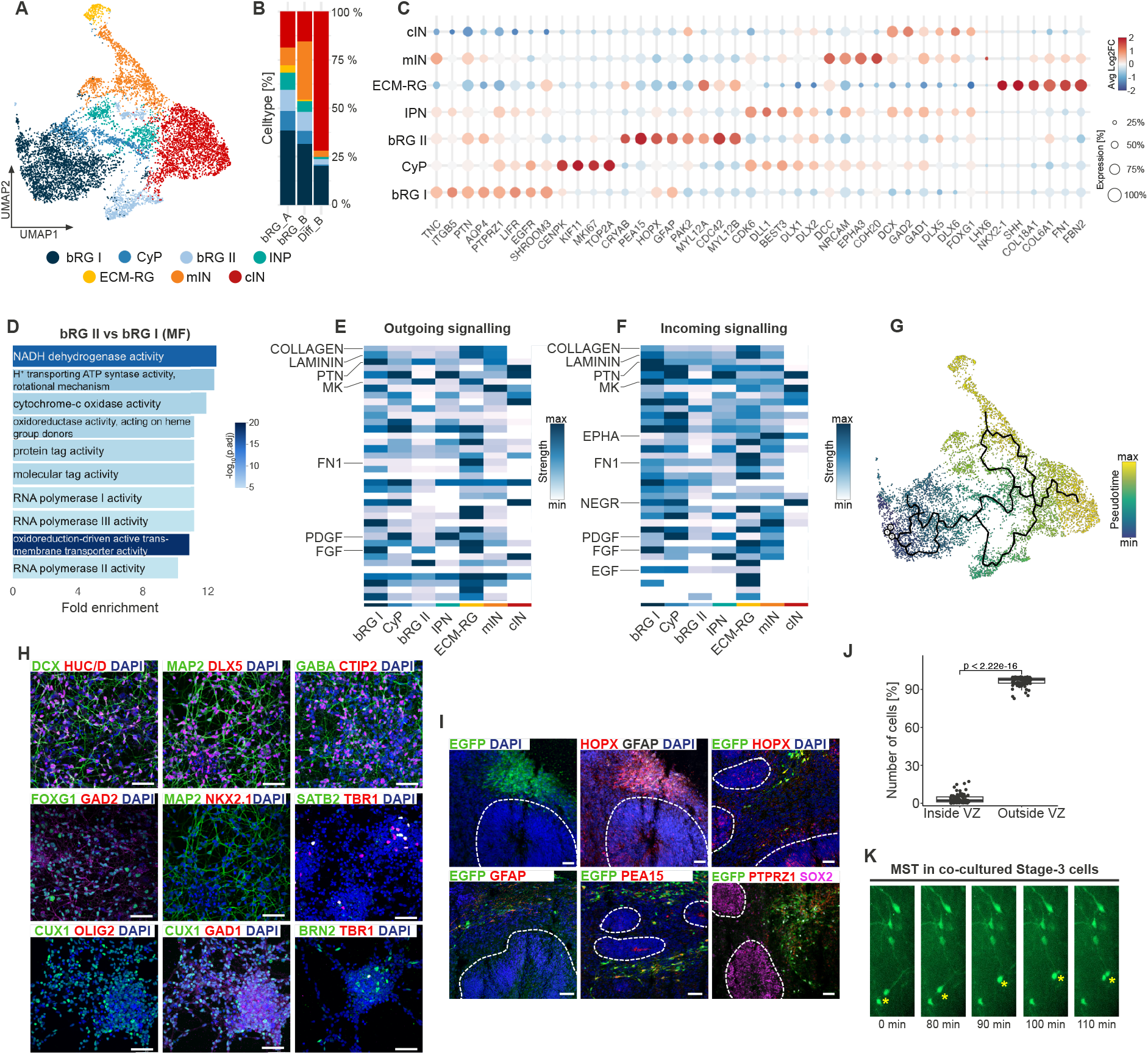
Cellular heterogeneity, lineage trajectories and functional integration of Stage-3 bRG. **A**. UMAP visualization of single-nuclei RNA sequencing data showing identified cell populations. **B**. Cell type distribution across samples. **C**. Dot plot of marker gene expression used for cluster annotation. **D**. GO enrichment (molecular function) comparing bRG-II versus bRG-I. **E-F** Predicted outgoing and incoming signalling interactions across clusters **G**. Pseudotime trajectory analysis. **H**. Immunostaining of Stage-3-derived neurons. **I**. Engraftment of EGFP-labelled Stage-3 bRG into organoid slices. **J**. Quantification of EGFP+ cells localization inside versus outside ventricular domains. **K**. MST in engrafted bRG. Scale bars: 50 µm.

To assess functional competence in a complex environment, EGFP-labeled Stage-3 bRG were seeded onto forebrain organoid slices. EGFP^+^ cells localized predominantly outside VZ– organized regions, with only ∼2% detected within VZ domains (**Fig. 2I-J**). Engrafted cells retained expression of RG and bRG-associated markers (HOPX, PTPRZ1, GFAP, PEA15) and EGFP^+^ cells undergoing MST were observed within the organoid context (**Fig. 2I-K**).

This defined 2D system provides a scalable and controllable platform to interrogate human bRG biology across independent iPSC backgrounds. In contrast to organoid models, the reductionist 2D design enables precise manipulation of niche signals and systematic perturbation of bRG behaviours. Identification of PAK2 as a regulator of MST illustrates the platform’s capacity to uncover molecular mechanisms controlling bRG dynamics. By complementing organoid and fetal tissue models, this expandable 2D system provides a versatile platform to investigate human-specific aspects of corticogenesis and their dysregulation in NDDs and cancer.

## Acknowledgements

We thank Gina Tillmann and Helene Schamber for the pivotal technical support. We thank the DKFZ Single-Cell Open Lab (scOpenLab) for assistance with the scRNA sequencing experiments. We acknowledge the support of the High Throughput Sequencing Unit of the Genomics & Proteomics Core Facility of the German Cancer Research Center (DKFZ).

## Declarations

### Data and code availability

Raw data is available through the European Genome-Phenome Archive xxxx. All original code has been deposited at GitHub repository and is publicly available as of the date of publication (https://github.com/xxxx/xxxx).

### Author Contributions

Methodology: A.A., M.G., A.C.R., R.W., A.D.B., A.J., J.L.; Validation: A.A., M.G., R.W., Y.H; Investigation: A.A., M.G., A.C.R., Y.H., R.W., F.M., A.J., C.G.; Formal Analysis: A.A., M.G., A.C.R., R.P.F, A.H., R.W., J.L.; Seq Data analysis: A.A., M.G., A.H., R.P.F., J.L.; Visualization: A.A., M.G. J.L.; Writing – Original Draft: A.A., M.G., J.L.; Writing – Review & Editing: all authors; Conceptualization: P.K., A.D.B., A.J., J.L.; Supervision: P.K., A.D.B., A.J., J.L.; Funding Acquisition: P.K., A.D.B., J.L.; Project Administration: J.L.

---

**Supplementary Figure 1.**
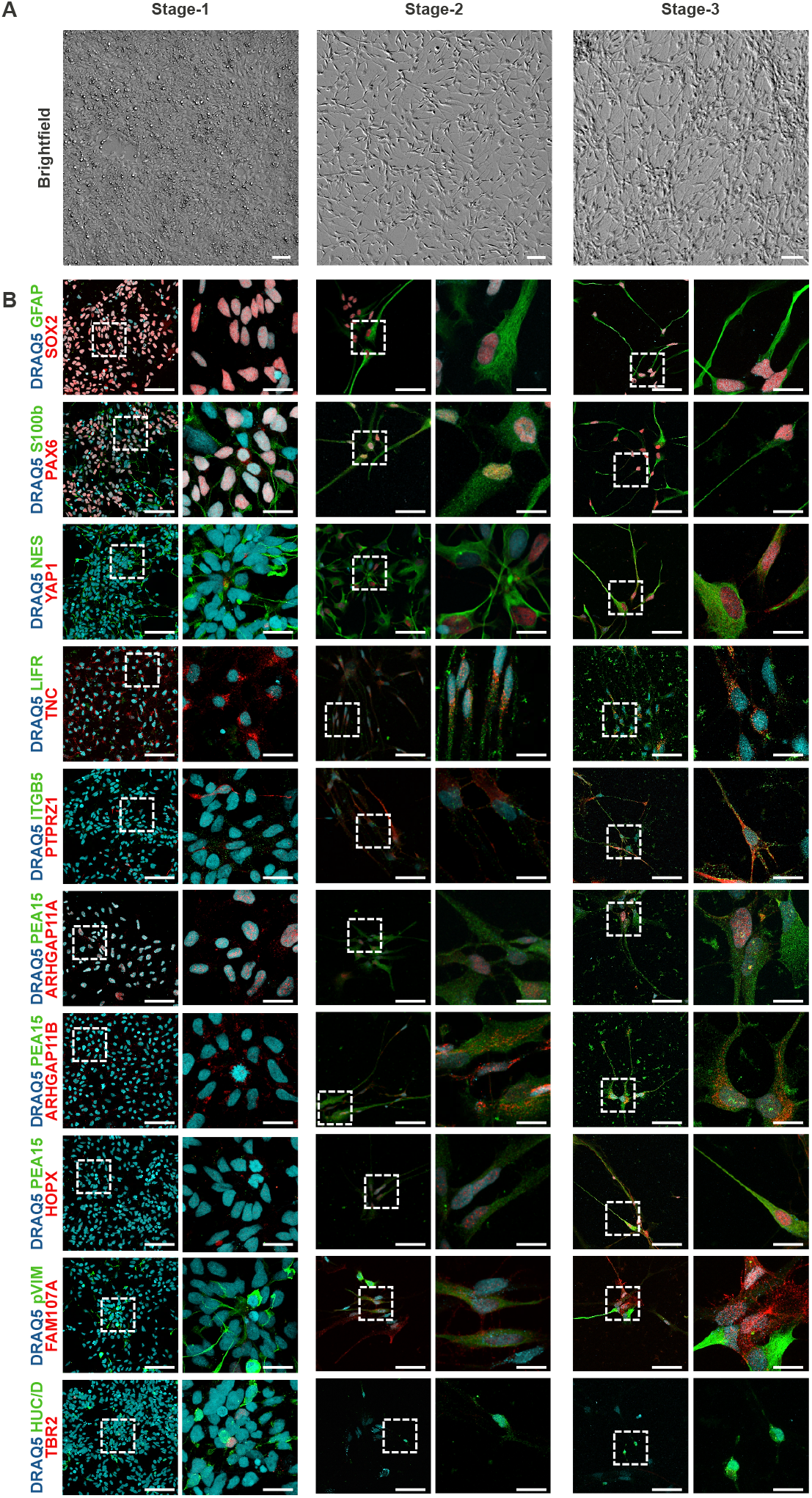
Validation of bRG differentiation across stages. **A**. Representative brightfield images illustrating morphological progression from iPSCs through Stage-1 and Stage-2 to Stage-3 cultures. Scale bars, 50 µm. **B**. Representative immunostainings of iPSC-derived cells at Stage-1, Stage-2 and Stage-3 stained for neural progenitor markers (SOX2, PAX6, NES, ARHGAP11A), glial-associated markers (GFAP, S100b, PEA15), RG niche / ECM components (TNC, LIFR), bRG-associated markers (YAP1, ITGB5, PTPRZ, HOPX, FAM11A, ARHGAP11B), intermediate progenitor marker (TBR2) and neuronal marker (HUC/D). Of note: nuclear localization of YAP1 is observed in a subset of Stage-2 cells and becomes more prominent in Stage-3 cultures. Similarly, ITGB5, PTPRZ1, HOPX and FAM107A stainings appear more widespread in Stage-3 compared to Stage-2 cultures. Insets show higher magnification views of individual cells. Nuclei were counterstained with DRAQ5. Scale bars, 50 µm (overview) and 10 µm (insets).

**Supplementary Figure 2.**
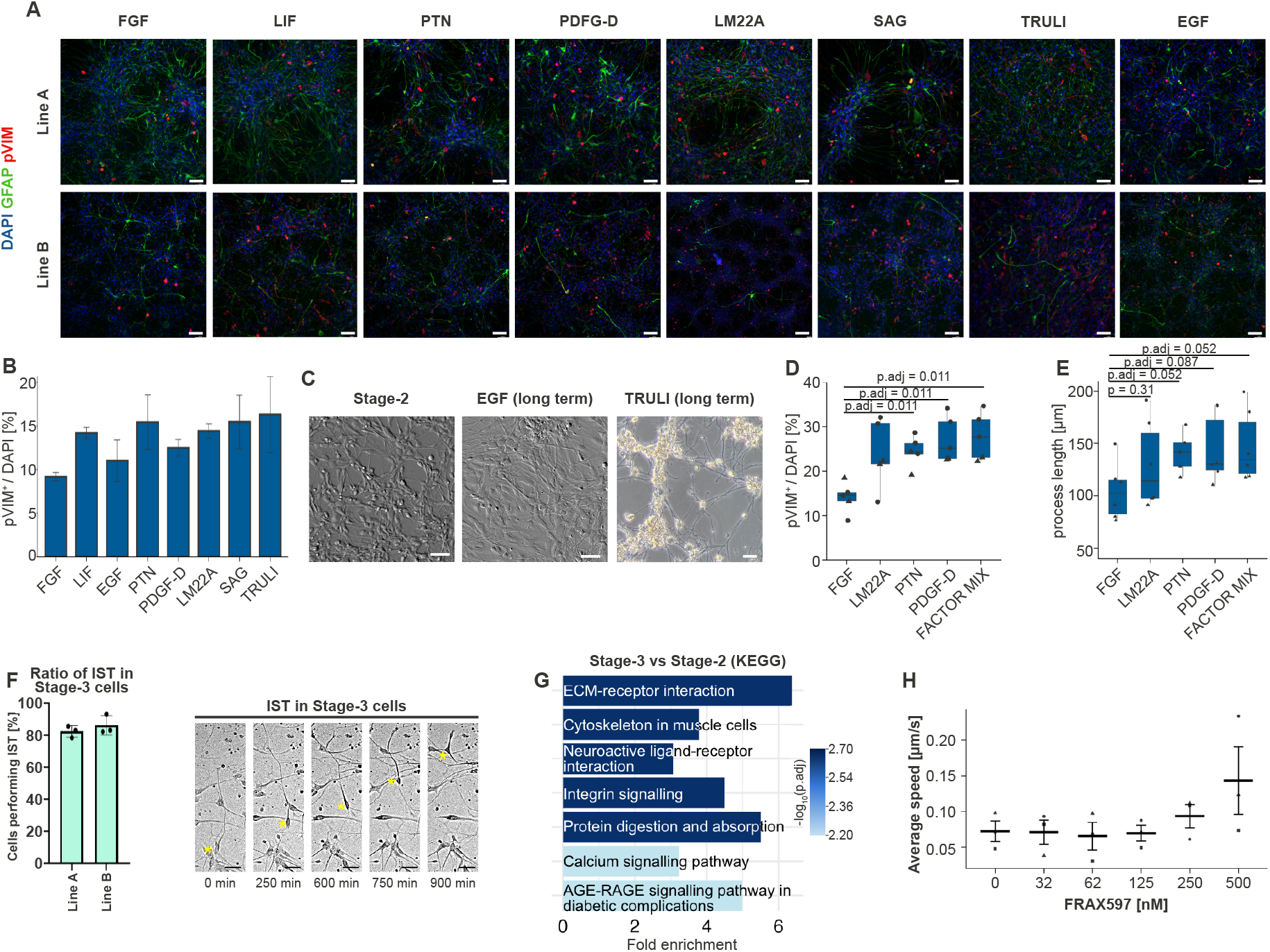
Screening, optimization and functional characterization of the niche factors promoting stabilization of bRG cultures. **A**. Representative immunostainings of Stage-2 cultures treated with candidate niche factors in addition to FGF2. Cells were stained for phosphorylated vimentin (pVIM) to label mitotic progenitors and GFAP to visualize RG-like morphology. Images are shown for two independent iPSC lines. Scale bars, 50 µm. **B**. Quantification of proliferative cells (pVIM^+^) in Stage-2 cultures treated with candidate niche factors during the initial niche-factor screening. Data represent two batches from two independent iPSC lines (327,413 total cells quantified). **C**. Representative brightfield images illustrating the effects of selected pathway modulators on Stage-2 RG cultures. Stage-2 RG cultures maintained with FGF2 alone, or supplemented for 3 weeks with EGF or the YAP activator TRULI. Both conditions resulted in altered culture morphology and reduced long-term stability. Scale bars, 50 µm. **D–E**. Quantification of proliferative cells (D; pVIM+ fraction) and average Nestin+ process length (E) in Stage-2 cultures treated with individual candidate factors or the final factor combination. Each point represents the mean value of one biological replicate. Circles and triangles indicate the two independent iPSC lines used. Data were obtained across five (pVIM) or six (Nestin) independent differentiation batches. Boxplots display the median (centre line) and interquartile range (box; 25th–75th percentiles), with whiskers extending to 1.5× the interquartile range. Statistical comparisons were performed on replicate means using Wilcoxon rank-sum tests against the FGF condition, with Benjamini–Hochberg correction applied for multiple comparisons. **F**. Quantification and representative time-lapse imaging of IST in Stage-3 bRG cultures. Each dot represents the mean fraction of cells undergoing IST per independent differentiation batch (2 lines, 3 batches each). Data are shown as mean ± SD. Representative time-lapse images illustrate IST progression at the indicated time points. Yellow asterisks mark cells undergoing IST. **G**. KEGG pathway enrichment analysis comparing Stage-3 and Stage-2 cultures, highlighting pathways associated with extracellular matrix interactions, integrin signaling, cytoskeletal regulation and neuroactive ligand-receptor signaling. **H**. Quantification of average speed in Stage-3 bRG cultures following treatment with different concentrations of the PAK2 inhibitor FRAX597. Speed was assessed during 96 h live-cell imaging with 10 min acquisition intervals. For each independent replicate (n=3), the mean speed was calculated across all tracked cells. Each point represents one independent replicate, defined as a unique combination of cell line and experimental batch. Data were obtained from two independent cell lines across three independent differentiation batches. Data is reported as mean ± SD. Statistical comparisons against the control condition were performed on replicate means using paired Wilcoxon signed-rank tests, with Benjamini-Hochberg correction applied for multiple comparisons, only p.adj < 0.05 are reported.

**Supplementary Figure 3.**
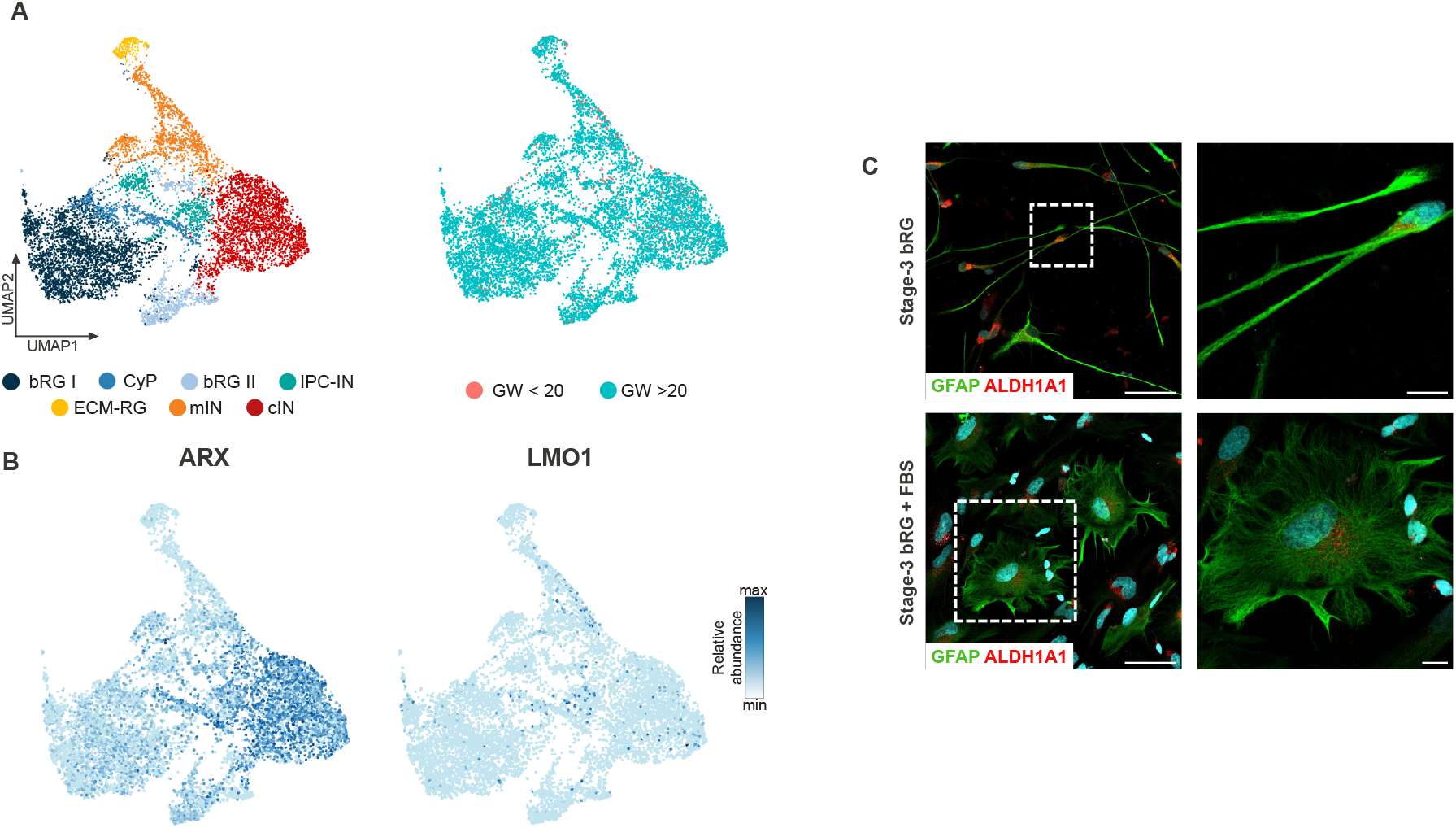
Transcriptional identity, lineage potential and reproducibility of Stage-3 bRG cultures. **A**. UMAP visualization of Stage-3 single-nuclei RNA sequencing data showing identified cell populations (left; reproduced from main Fig. 2 for direct comparison) and label transfer mapping onto a reference atlas of the developing human cortex^23^ (right), indicating predominant correspondence with post-GW20 developmental stages. **B**. Feature plots illustrating expression of the cortical interneuron-associated marker ARX within the cIN cluster, while the ganglionic eminence interneuron-associated marker LMO1 remain largely absent. **C**. Representative images illustrating astroglial differentiation of Stage-3 cultures. Top: Stage-3 bRG cultures maintained under standard conditions. Bottom: Cultures exposed to astroglial differentiation conditions showing GFAP^+^ cells with astroglia-like morphology. Dashed white boxes indicate regions shown at higher magnification on the right. Scale bars, 50 µm.

## Methods

### Culture of Human iPSC

Human iPSC lines used in this study were obtained from previously established sources. iPSC line A (44-year-old female donor, derived from skin fibroblasts) and iPSC line C (25 years old female donor, derived from skin fibroblasts) were provided by Dr. Sandra Horschitz (Central institute of Mental Health) under approval of the Ethics Committee II of Medical Faculty Mannheim of Heidelberg (approval no. 2014-626N-MA). iPSC cell line B (2-years old female donor) was derived from skin fibroblasts obtained from Coriell Biorepository (catalog ID GM00969). All participants provided written informed consent for the derivation and use of iPSCs in research.

iPSCs were maintained as colonies on Geltrex-coated (GT, Thermo Fisher Scientific) cell culture plates in Essential 8 (E8) medium supplemented with 10 U/ml penicillin and 10 µg/ml streptomycin (Thermo Fisher Scientific), with daily medium change. Cells were split using 500 µM EDTA (Thermo Fisher Scientific) at a 1:3 to 1:10 ratio. After passaging, the medium was supplemented with 5 μM Y-27632 (Cell Guidance Systems) to enhance survival. All lines were routinely tested and confirmed negative for mycoplasma. Pluripotency and genomic integrity were previously validated as described^26^.

### Generation of iPSC-derived bRG

Differentiation into Stage-1 NE cells was adapted from a previously established cortical induction protocol^26^. Briefly, iPSC were grown to 95-100% confluence before initiation of neural induction. Neural induction was performed in neural induction medium, consisting of base medium composted of DMEM/F12, (Thermo Fisher Scientific) supplemented with 1% v/v GlutaMAX (Thermo Fisher Scientific), 1% v/v non-essential amino acids (NEAA, Thermo Fisher Scientific), 50 µg/ml gentamicin (Thermo Fisher Scientific), 0.5% v/v N2 supplement, 0.8 mg/ml D-glucose (Sigma Aldrich), and 50 µM 2-mercaptoethanol (Thermo Fisher Scientific). For neural induction, the base medium was additionally supplemented with 1% v/v B27 (Thermo Fisher Scientific) and the small-molecule SMAD inhibitors LDN-193189 (0.2 μM, Milteny), A83-01 (0.5 μM Tocris), and the canonical WNT inhibitor XAV939 (2 μM, Cell Guidance Systems). Cells were maintained under neural induction medium for 7-10 days. Cultures were then dissociated using TrypLE (Thermo Fisher Scientific) or Accutase (BioLegend) and replated at a 1:2 ratio onto GT-coated culture plates (1:100 dilution) in neural induction medium. Subsequently, cells were passaged 1-2 additional times at a 1:2 ratio every 3-4 days in neural induction medium. Cultures maintained under these conditions for up to three passages were defined as Stage-1 NE cells.

Stage-1 NE cultures were further processed following an approach adapted from the protocol described by Ostermann et al^11^. To allow spontaneous differentiation and depletion of proliferative Stage-1 NE cells, cultures were maintained for 5 weeks in base medium supplemented with 1% B27 v/v without exogenous growth factors, with daily medium changes. Following this differentiation period, cultures were dissociated using TrypLE and replated as single cells at high density onto Geltrex-coated culture plates. To selectively expand growth factor–responsive neural stem/progenitor cells, cultures were subsequently maintained in base medium supplemented with FGF2 (10 ng/ml, Qkine) for approximately 4 weeks, with regular passaging. Cultures maintained under these conditions were defined as Stage-2 RG.

To stabilize and enrich the bRG state, Stage-2 cultures were transitioned to base medium supplemented with the growth/niche factors FGF2 (10 ng/ml), PTN (25 ng/ml, PreproTec), and PDGF-D (20 ng/ml, R&D systems). The neurotrophic TrkB agonist LM22A (1 µM, Sigma Aldrich) was included to support cell survival and growth. Cultures were maintained under these conditions for at least 2 weeks, after which they were defined as Stage-3 bRG. During optimization of the culture conditions, additional growth factors and pathway modulators were tested, including EGF (20 ng/ml, Qkine), SHH agonist SAG (1 μM, Adipogen), LIF (20 ng/ml, Qkine) and the YAP activator TRULI (1 μM, AmBeed). These conditions were not included in the final medium due to reduced long-term culture stability or potential effects on regional identity or lineage output.

Stage-2 and Stage-3 cultures were routinely passed upon reaching confluency or as required for experimental procedures using TrypLE or Accutase and replated onto GT-coated cell culture plates (1:100 dilution). Stage-2 cultures were typically maintained at high (∼90 %) confluency during routine expansion, whereas Stage-3 cultures tolerated both high- and low (∼20 %)-density culture conditions. Stage-3 cultures could be expanded for multiple passages; however, experiments were typically performed within the first ∼15 passages to minimize the risk of long-term culture–associated genomic alterations. Cells at Stage-1, Stage-2 and Stage-3 were cryopreserved in freezing medium consisting of base medium supplemented with 10% v/v DMSO (Sigma Aldrich) and 20 μM Y-27632 (Cell Guidance Systems).

To assess robustness of the protocol, the differentiation strategy was applied across multiple independent iPSC lines in addition to the lines presented in this study. The protocol consistently generated expandable bRG-like cultures across these genetic backgrounds. Experiments were not randomized, and investigators were not blinded to differentiation conditions. Sample sizes were not predetermined by statistical methods.

### Neuronal differentiation of bRG

For spontaneous differentiation (used for single-nuclei RNA sequencing), confluent Stage-3 bRG cultures were maintained in base medium supplemented with 1% v/v B27 without additional growth factors. Cultures were maintained under these conditions for up to three months, with regular medium changes, allowing gradual neuronal differentiation. For directed neuronal differentiation (used for immunocytochemical validation), confluent Stage-3 bRG were switched to Neurobasal medium (Thermo Fisher Scientific) supplemented with 1% v/v N2 supplement, 2% v/v B27 (without vitamin A), 1% v/v NEAA, 1% v/v Glutamax, 50 µg/ml gentamicin, 1 µM LM22A, 5µM DAPT (Cell Guidance Systems), and 2.5 µg/ml insulin (Sigma Aldrich). Cultures were maintained under these conditions for two weeks, with regular medium changes, before fixation and downstream analysis

### Astrocyte differentiation of bRG

For directed astrocyte differentiation, confluent Stage-3 bRG cultures were maintained in base medium supplemented with 10% fetal bovine serum (FBS, Thermo Fisher Scientific). Cultures were maintained under these conditions for 2 weeks with regular medium changes before fixation and downstream analysis.

### Generation of forebrain organoids

Forebrain organoids were generated as previously described^26,27^. Briefly, dissociated iPSCs were seeded in low attachment U-bottom 96-well plates in E8 medium supplemented with 50 µM Y-27632, 10 U/ml penicillin and 10 µg/ml streptomycin. Neural induction was initiated at day 5, using dual-SMAD and WNT inhibition (0.2 μM LDN-193189, 0.5 μM A83-01, 2 μM XAV939). Upon formation of translucent neuroectoderm (day 9-11), aggregates were embedded in GT and cultured in differentiation medium under continuous orbital agitation (70 rpm). From day 40 onwards, organoids were maintained in maturation medium supplemented with 1% v/v B27, 1 µM LM22A, 2 µM LM22B, 10 ng/ml GDNF, 200 nM ascorbic acid and 0.2 % v/v GT. Between day 35-40, organoids were embedded in 2% w/v low-melting agarose (Sigma Aldrich) and sectioned into 300 µm slices using a vibratome. Organoid slices were subsequently maintained in maturation medium under continuous orbital agitation (70 rpm) until further use.

### Coculture of Stage-3 bRG with forebrain organoid slices

EGFP-labeled Stage-3 bRG were generated by lentiviral transduction (pLentiPGK-EGFP-SV40-blasticidine). Transduced cells were selected with 10 μg/ml blasticidin (Carl Roth) for ≥2 weeks and homogenous EGFP expression was confirmed by fluorescence microscopy. For coculture, 100,000 EGFP-labeled Stage-3 bRG were seeded onto 300 µm forebrain organoid slices (day 40-50) in maturation medium supplemented with FGF-2 in a 48-well plate (one slice per well). FGF-2 was removed after 24 h to facilitate infiltration. Cultures were briefly maintained under orbital agitation to remove non-integrated cells, then transferred to air-liquid interface membranes and maintained for 10 days before fixation. For coculture quantification, VZ regions were segmented using Imaris “surface detection” and EGFP+ cells were quantified using “Spots” detection.

### Immunofluorescence

Cells were fixed in 4% paraformaldehyde (PFA, Sigma Aldrich) for 10 min and processed for immunostaining or stored at 4°C. Organoids were fixed, embedded, and cryosectioned (20 μm) as previously described^26^. Samples were blocked and permeabilized for 1 h at room temperature (RT) in PBS containing 10% fetal bovine serum (Thermo Fisher Scientific), with or without 0.3%, Triton X-100 (Sigma Aldrich), incubated with primary antibodies (**Supp. Table 12**) at 4°C overnight (16 h), washed three times with PBS, and incubated with secondary antibodies for 1 h at RT. Nuclei were counterstained with DAPI and samples were mounted in Mowiol. Images were acquired using Leica TCS SP5II or Stellaris 5 confocal microscopes, or a Leica DM6 B fluorescence microscope, and processed with Leica Application Suite AF, Leica Application Suite X, and ImageJ.

### MST and IST quantification

For live imaging experiments, Stage-2 RG and Stage-3 bRG were plated at ∼30% confluence on GT-coated plates to enable single-cell tracking. Live imaging of MST events was performed for 96 h using a CellDiscoverer 7 platform (Zeiss), acquiring images every 10 min at ≥3 positions per well. Dividing cells were detected using Imaris “Spot” tracking based on transient cell rounding. MST events were defined as mitoses in which the soma displaced by >1 standard deviation from its initial position along a linear trajectory. For pharmacological inhibition, FRAX597 (MedChemExpress) was added 24 h prior to imaging at the indicated concentrations. Mitosis rate was calculated as the total number of mitotic events normalized to the number of cells present at time 0. For live imaging of IST events, bRG cells were plated onto 6-well glass bottom plates (IBL P06-1.5H-N), GT-coated and imaged on a Nikon TI2-E inverted video-microscope equipped with a photometric Kinetix sCMOS camera, and each position was imaged every 5 minutes in phase contract (DIC) with a 10X Plan Fluor, NA: 0.20, WD 15.20 dry objective, using a perfect focus (PFS) mode. All experiments were performed in triplicate across two genetic backgrounds unless otherwise stated.

### Culture of primary bRG

Human fetal tissue samples were collected by the laboratory of Alexandre Baffet (Agence de la Biomédecine, approval number: PFS17-003) with prior patient consent and in accordance with legal and institutional ethical regulations. Primary human bRG were isolated and cultured by the Baffet laboratory as previously described (Wimmer et al., 2026). In brief, fresh human fetal prefrontal cortex tissue was mechanically dissociated and plated onto poly-D-lysine and fibronectin-coated culture dishes in DMEM/F12-based medium supplemented with B27 (−vitamin A), FGF (20 ng/mL), and EGF (20 ng/mL). Media was replaced the following day and every two days thereafter to remove non-adherent cells and debris, enriching for radial glial populations. After 2–3 weeks in culture, cells were passaged using Accutase and replated onto coated dishes. bRG identity was confirmed by immunostaining for established marker expression and maintained under these conditions. For comparative analyses, cells were subsequently transferred to the culture conditions used for iPSC-derived bRG and maintained for 2 weeks prior to sequencing experiments.

### Bulk RNA sequencing and analyses

For bulk RNA sequencing, RNA was extracted from three independent biological replicates per differentiation stage for iPSC-derived cultures (Stage-1, Stage-2, and Stage-3), corresponding to three independent differentiations of the same iPSC cell line. For primary bRG, samples were obtained from three developmental stages (GW14, GW17, and GW21). For each stage, one preparation was expanded and RNA was collected from four wells, which were treated as technical replicates. RNA was extracted using TriFast (VWR) according to the manufacturer’s protocol. RNA integrity was assessed using an Agilent Bioanalyzer 2100 (RNA 6000 Nano Kit), and samples with RIN > 7.5 were used for library preparation. Libraries were prepared using the TruSeq Stranded protocol (Illumina) and sequenced (single-end, 50 bp) on a NovaSeq 6000 platform. Reads were quantified using featureCounts, and count matrices were imported into R for downstream analysis using DESeq2^28^. Genes with zero total counts across all samples were removed prior to model fitting. Differential expression analysis was performed using a design formula including batch and condition. Size factors, dispersion estimates, and model coefficients were calculated using the DESeq function. Pairwise contrasts were computed using the results function, applying a Benjamini–Hochberg adjusted p-value threshold of ≤ 0.05. Differentially expressed genes (DEGs) were defined by adjusted p ≤ 0.05. Volcano plots were generated using log_2_ fold change and −log10 adjusted p-values. Genes belonging to a curated bRG marker list were highlighted where indicated.

#### Functional enrichment analysis

For selected contrasts, significantly regulated genes were subjected to Gene Ontology (GO) enrichment analysis using clusterProfiler (enrichGO)^29^. Gene symbols were converted to Entrez identifiers using org.Hs.eg.db. Enrichment analyses were performed separately for Biological Process (BP), Molecular Function (MF), and Cellular Component (CC) ontologies. Terms with adjusted p-value < 0.05 (Benjamini–Hochberg correction) were considered significant. Results were exported as supplementary tables and visualized as bar plots. All analyses were conducted in R using DESeq2, clusterProfiler, and associated packages. Scripts and session information are provided.

### MST-associated protein-protein interaction (PPI) network inference

Genes significantly upregulated in Stage-3 versus Stage-2 cells (log2FC ≥ 0.5, adjusted p. ≤ 0.05) were filtered to protein-coding genes with annotated localization to the cytoskeleton or plasma membrane. To assess potential association with MST, a literature mining strategy was applied. PubMed searches were performed using predefined MST-related search terms (**Supp. Table S10**), and abstracts were screened for predefined keywords (**Supp. Table S11**). Keyword counts were adjusted using a skewed Gaussian weighting function (µ = average_term_count, σ = sd_term_count, a = -2) to increase the relative contribution of low-frequency, specific terms. Scores were normalized to the theoretical maximum to allow comparability across genes. Log2FC were incorporated using a second skewed Gaussian weighting (µ = 4, σ = 1.2, a = -1.5) to reduce overrepresentation of highly upregulated genes. Final composite scores were calculated as the product of normalized literature scores and weighted log2FC values. Genes were ranked accordingly. The top 25% of ranked genes were used as seed notes to construct PPI networks using STRINGv11.5. First-shell interactors were retrieved and filtered for expression in the Stage-3 dataset (upregulated or not significantly downregulated genes). To reduce network complexity, nodes located more than three edges from any seed node were removed and only the largest connected component was retained. Edges were weighted using a composition metric integration: (i) STRING interaction confidence (weight 0.75), (ii) PubMed-derived association score (weight 0.5), and (iii) gene expression score (weight 0.25). Edge scores were calculated as the sum of the expression scores of connected seed nodes. An empirical cutoff of 450 was applied to remove low-confidence interactions. Network clustering was performed in Cytoscape v3.10.3^30^ using the Markov Cluster algorithm (MCL; inflation 2.4). Eigenvector centrality was computed using CytoNCA to identify highly connected regulatory candidates.

### Single nuclei RNA sequencing (snRNA-seq)

Stage-3 bRG cultures derived from two independent iPSC lines (line A and B), as well as their corresponding differentiated neuronal populations, were processed for single-nuclei RNA sequencing. Cultures were maintained in expansion conditions without passaging for extended periods (≥4 weeks) prior to nuclei isolation. Neuronal differentiation was induced by growth factor withdrawal, as described above. The library from differentiated neurons of line A was excluded due to unsuccessful sample demultiplexing, preventing reliable assignment of reads. Cells were dissociated using TrypLE Express (Thermo Fisher Scientific) and pelleted by centrifugation (800 × g, 4 min, room temperature). Nuclei were isolated by mechanical disruption in TST buffer (146 mM NaCl, 10 mM Tris-HCl pH 7.5, 1 mM CaCl_2_, 21 mM MgCl_2_, 0.03% Tween-20, 0.01% BSA), followed by filtration through a 30 µm strainer. After centrifugation (500 × g, 5 min, 4°C), nuclei pellets were resuspended in ST buffer (292 mM NaCl, 20 mM Tris-HCl pH 7.5, 2 mM CaCl_2_, 42 mM MgCl_2_) supplemented with RNase inhibitor (1 U/µl) and filtered through a 20 µm strainer. Nuclei concentration and quality were assessed prior to library preparation. For multiplexing, nuclei were hashed using oligonucleotide barcodes: 500,000 nuclei per sample were incubated with the corresponding oligo at a final concentration of 1.8 µM for 20 min on ice. Nuclei were then washed in PBS supplemented with 1% (w/v) BSA and 0.2 U/µl RNase inhibitor, followed by centrifugation (400 × g, 10 min, 4°C). This wash step was repeated three additional times. Nuclei were subsequently resuspended in the final buffer, counted, and quality-controlled prior to pooling for library preparation. Gene expression libraries were generated using the Chromium Single Cell 3’ v3.1 chemistry with Feature Barcoding (10x Genomics) according to the manufacturer’s instructions. Libraries were processed by the High-Throughput Sequencing Unit of the Genomics and Proteomics Core Facility at the German Cancer Research Center (DKFZ) and sequenced on an Illumina NovaSeq 6000 platform (S1 flow cell, paired-end 28 × 94 bp).

### snRNA-seq data quality control and analysis

FASTQ files were processed using Cell Ranger (10x Genomics) to generate filtered feature barcode matrices. Downstream analyses were performed in R using Seurat v5.0.266, unless otherwise stated^31^. Filtered matrices were imported using Read10X and converted into Seurat objects. For multiplexed samples, hashtag oligonucleotide (HTO) counts were processed separately, normalized using centered log-ratio (CLR) transformation, and demultiplexed using HTODemux (positive quantile threshold = 0.995). Only singlet nuclei were retained. Quality control filtering excluded nuclei with <1,350 or >9,000 detected genes and those with ≥10% mitochondrial transcripts. Data were log-normalized and 2,000 variable features were identified using the variance stabilizing transformation (VST) method. Cell cycle phase scores (S and G2/M phase) were calculated using canonical gene sets (Seurat::cc.genes), and percent mitochondrial content together with S.Score and G2M.Score were regressed out during scaling.

#### Dataset integration, clustering and annotation

Independent datasets were integrated using Seurat’s anchor-based integration workflow. Integration anchors were identified using the first 30 principal components, and data were integrated (dims = 1:30). The integrated assay was used for downstream analyses. Principal component analysis (PCA) was performed on the integrated assay. Based on elbow and variance-explained analyses, the first 12 principal components were retained. UMAP dimensionality reduction was computed using these components. A shared nearest neighbor (SNN) graph was constructed using FindNeighbors (dims = 1:12), and clustering was performed using the Leiden algorithm (resolution = 0.35). Clusters were annotated manually using established marker genes and a curated list of bRG markers.

#### Differential expression and enrichment analysis

Differential expression analyses were performed using FindMarkers with a minimum detection difference of 0.3. Genes with adjusted p-value < 0.05 were retained. Ribosomal protein genes (RPL/RPS) were excluded from enrichment analyses. Significantly upregulated genes were converted to Entrez identifiers and subjected to Gene Ontology (GO) enrichment analysis using clusterProfiler (enrichGO) across Biological Process, Molecular Function, and Cellular Component ontologies. Multiple testing correction was performed using the Benjamini–Hochberg method.

#### Trajectory interference

Trajectory analysis was performed using Monocle3^32^. The integrated UMAP embedding was transferred to a Monocle3 cell data set. Cells were clustered, a trajectory graph was learned without partitioning, and pseudotime was inferred using default parameters.

#### Cell-cell communication analysis

Cell–cell communication was inferred using CellChat^33^ with the human ligand–receptor database. Normalized expression data were grouped by annotated cell type. Overexpressed genes and interactions were identified, communication probabilities were computed, and signaling pathways were aggregated. Interactions were filtered to retain pathways represented by at least 10 cells per group. Outgoing and incoming signaling centrality were quantified using network centrality analysis and visualized as pathway-level heatmaps.

All analyses were performed in R. Figures were generated using Seurat, ggplot2, and associated packages. Session information and scripts are provided as supplementary information for this manuscript.

### Statistics

Quantification of immunofluorescence and live-cell imaging data was performed in R (v4.4.2). For all assays, the experimental unit was defined as a biological replicate corresponding to a unique combination of cell line and differentiation batch. Images or single-cell measurements obtained within the same replicate were treated as technical subsamples and averaged prior to statistical testing. Statistical inference was therefore performed on replicate-level values. For immunostaining quantification of phosphorylated vimentin (pVIM), the fraction of positive cells was calculated for each field as the number of pVIM-positive cells divided by the number of DAPI-positive nuclei and expressed as a percentage. Values were averaged across fields within each biological replicate (cell line × batch). In total, five independent differentiation batches from two cell lines were analyzed. The number of cells analyzed per condition was: FGF (n = 2,894), MIX (n = 4,683), LM22A (n = 3,238), PTN (n = 2,756) and PDGFD (n = 4,152). For neurite process length measurements, the mean process length was calculated across all measured cells within each replicate. In total, six independent differentiation batches from two cell lines were analyzed. The number of cells analyzed per condition was: FGF (n = 815), MIX (n = 806), LM22A (n = 841), PTN (n = 794) and PDGFD (n = 653). For live-cell imaging assays of cell motility, individual cell trajectories were used to compute instantaneous speeds, and the mean speed across all tracked cells within each replicate was calculated. In total, six independent differentiation batches from two cell lines were analyzed. The number of cells analyzed per condition was: 0 nM (n = 788), 31.25 nM (n = 406), 62.5 nM (n = 546), 125 nM (n = 574), 250 nM (n = 1,043) and 500 nM (n = 1,066). For mitotic somal translocation (MST) analyses, MST frequency was computed for each replicate as the proportion of MST events relative to the total number of mitotic events. Comparisons between two matched experimental conditions were performed using Wilcoxon signed-rank tests on replicate means. For the analysis shown in Figure 1E, a one-tailed Wilcoxon signed-rank test (alternative = greater) was applied. In total, 3 independent differentiation batches from two cell lines were analyzed. The total number of mitosis per condition was: 3418 (Stage 2) 1662 (Stage 3). For experiments involving comparisons of multiple treatments against a reference condition, Wilcoxon rank-sum tests were performed on replicate means, and P values were adjusted for multiple testing using the Benjamini–Hochberg procedure where applicable. For the FRAX592 dose–response analysis of MST frequency, normalized MST values were modeled as a function of drug concentration including the control condition. To assess a monotonic concentration-dependent effect while accounting for matched experimental series, concentration was transformed as log10(dose + 1) and a linear model of the form normalized_MST ∼logDose + Paired was fitted, where Paired encoded matched replicate series. The significance of the dose-dependent trend was evaluated from the logDose term using the model summary and analysis-of-variance output. In all boxplots, the center line represents the median, the box indicates the interquartile range (25th–75th percentiles), and whiskers extend to 1.5× the interquartile range. Individual points correspond to replicate-level measurements. The total number of mitoses per concentration was: 0 nM (n = 1,167), 31.25 nM (n = 476), 62.5 nM (n = 402), 125 nM (n = 258), 250 nM (n = 81) and 500 nM (n = 10). All statistical tests were two-sided unless otherwise specified.

